# Periendothelial enrichment of PIEZO1 and its contribution to cisplatin tolerance in small cell lung cancer

**DOI:** 10.64898/2026.10.11.758385

**Authors:** Zhongliang Wang, Lingge Zeng, Xue Sun, Man Li, Rong Li, Fanrui Zeng

**Affiliations:** Department of Radiation Oncology, The First Affiliated Hospital of Guangzhou Medical University, Guangzhou Medical University, Guangzhou, 510000, China; School of Medical Sciences, Faculty of Medicine and Health, The University of Sydney, Camperdown, NSW 2006, Australia; Department of Pathology, Zhujiang Hospital, Southern Medical University, Guangzhou, 510282, China; Department of Medical Oncology, Shanghai Pulmonary Hospital, Tongji University Medical School Cancer Institute, Tongji University School of Medicine, Shanghai, 200433, China; Department of Pathology, Sichuan Provincial People’s Hospital, University of Electronic Science and Technology of China, Chengdu, 610072, China

**Keywords:** small cell lung cancer, PIEZO1, cisplatin, neuroendocrine state, spatial transcriptomics, YAP1

## Abstract

Small cell lung cancer (SCLC) contains neuroendocrine and non-neuroendocrine cell states that differ in their response to chemotherapy. We asked where cells expressing the mechanosensitive channel PIEZO1 reside in SCLC and how PIEZO1 affects cisplatin response. PIEZO1 expression increased in two resistant cell models and correlated inversely with neuroendocrine programmes in patient tumours. In a primary SCLC section profiled by single-cell spatial transcriptomics, 273 of 7,101 epithelial candidates expressed detectable PIEZO1. The odds of detection decreased with distance from annotated endothelium (adjusted odds ratio 0.58, 95% confidence interval 0.48–0.70 per doubling of [distance + 10 µm]). Within the same cells, a one-standard-deviation increase in neuroendocrine contrast corresponded to an odds ratio of 0.43 (0.37–0.48) after adjustment for endothelial distance and local composition. PIEZO1 knockdown lowered the cisplatin IC_50_ from 15.7 to 6.4 µg/mL in H69AR cells and from 15.7 to 5.9 µg/mL in H446DDP cells. In the parental lines, the PIEZO1 agonist Yoda1 increased the calcium-sensitive Fura-2 ratio, reduced cytoplasmic YAP1 and protected cells from cisplatin-induced apoptosis. Verteporfin partially reversed this protection. These findings localize a PIEZO1-expressing, low-neuroendocrine epithelial population near endothelium in human SCLC and identify a contribution of PIEZO1 to cisplatin tolerance in resistant cells.

## Introduction

Platinum-based chemotherapy produces an initial response in many patients with small cell lung cancer (SCLC), yet relapse is common. SCLC cells vary in their expression of neuroendocrine (NE) lineage programmes, and individual tumours can contain cells at different positions along this spectrum [1, 2, 3]. YAP-associated states expand in some relapsed tumours, and induction of YAP can alter cell identity and chemotherapy response in experimental models [4, 5]. How these states are organized within the tissue, and which signals they can use to survive platinum exposure, remain open questions.

PIEZO1 is a mechanically activated cation channel that couples membrane deformation to calcium entry [6]. Endothelial cells express PIEZO1, creating a specific problem for its study in tumours. A PIEZO1 signal near a vessel could arise from endothelial cells themselves, from nearby tumour cells or from both. Bulk expression and multicellular spatial spots do not resolve these possibilities. Here we combined tumour proteogenomics, single-cell atlases and spatial RNA measurements with genetic and pharmacological perturbation. We first identified the epithelial cells in which PIEZO1 is detected, then measured the contribution of PIEZO1 to cisplatin response in paired parental and resistant SCLC models.

## Results

### PIEZO1 expression rises in resistant SCLC models

We compared two SCLC cell-line pairs with different selection histories. H69AR is a doxorubicin-selected multidrug-resistant derivative of H69 [7], whereas H446DDP was selected for cisplatin resistance from H446 [8]. In H69AR, a joint RNA and N^6^-methyladenosine (m^6^A) screen nominated PIEZO1 among genes with higher RNA abundance and greater estimated immunoprecipitation enrichment (Fig. 1a). RNA sequencing identified 2,166 genes with increased expression and 2,103 with decreased expression relative to H69. ASCL1, INSM1, CHGA and SYP decreased, while REST, VIM, YAP1, WWTR1 and PIEZO1 increased (Fig. 1b,c). Epithelial-to-mesenchymal transition ranked first among enriched Hallmark programmes (normalized enrichment score 2.30), accompanied by hypoxia and Notch signalling (Fig. 1d,e). Library quality, programme tests and regional RNA coverage are shown in Fig. S1a–f.

**Figure 1.**
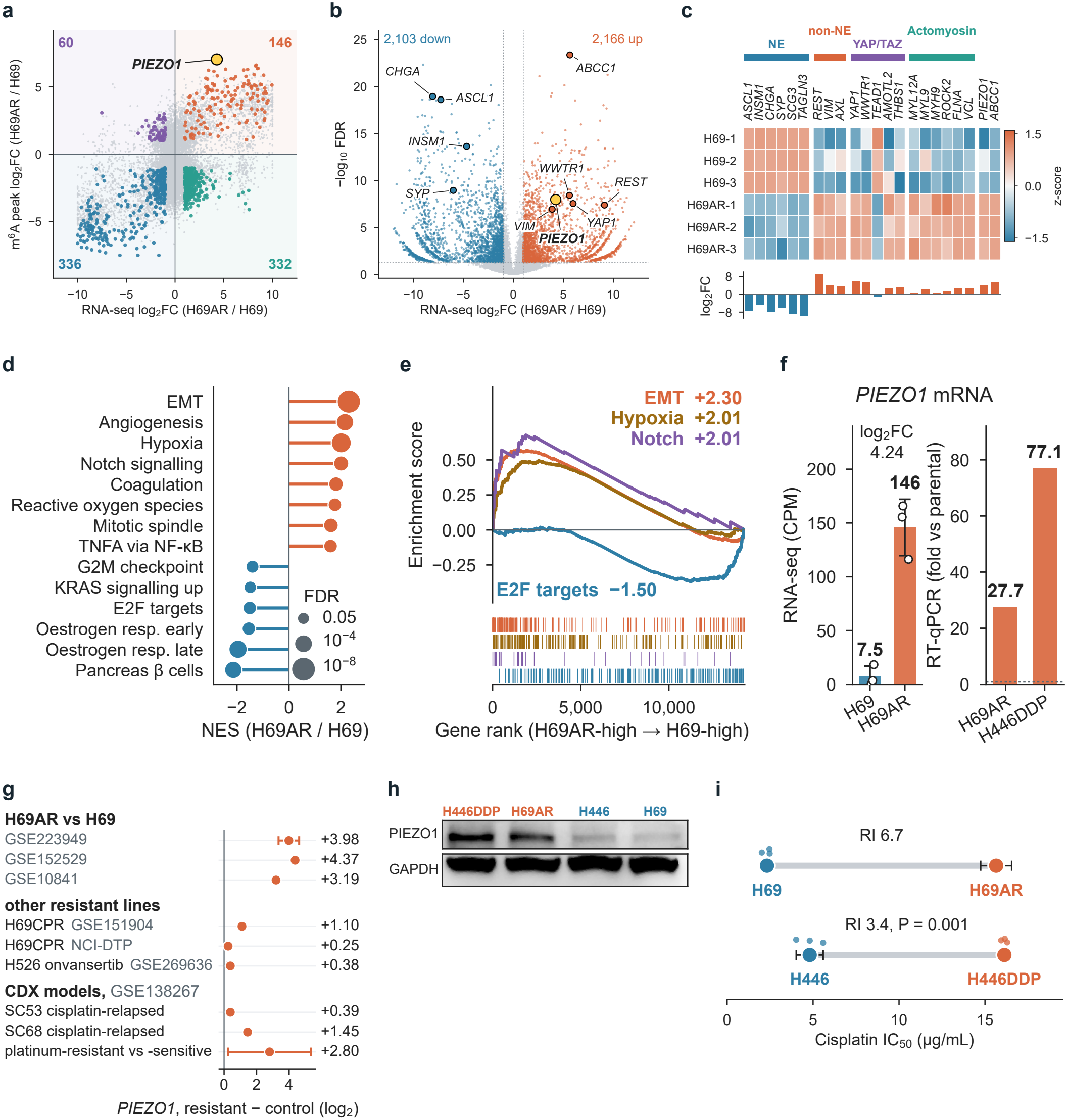
PIEZO1 expression in drug-resistant SCLC models. **a**, Differential enrichment in the m^6^A screen versus RNA abundance for H69AR relative to H69. **b**, Differential gene expression at FDR < 0.05 and absolute log_2_ fold change > 1. **c**, Expression of selected state, cytoskeletal and resistance genes across three libraries per line. **d**,**e**, Hallmark pathway enrichment and selected enrichment curves. **f**, PIEZO1 RNA by sequencing and the reported RT–qPCR fold change for each resistant–parental pair. **g**, Resistant-minus-control expression differences in nine public comparisons. Horizontal bars show 95% CIs where estimable. **h**, PIEZO1 immunoblots with GAPDH controls. **i**, Cisplatin IC_50_ and resistance indices in the H69/H69AR and H446/H446DDP pairs. The H69AR estimate is the siRNA control also used in Fig. 4b. The dispersion measure for that control was unspecified in the original record. Error bars for the other three estimates denote SD.

PIEZO1 RNA increased from 7.5 counts per million in H69 to 146 in H69AR. The RT–qPCR summaries showed 27.7-fold higher PIEZO1 expression in H69AR than in H69 and 77.1-fold higher expression in H446DDP than in H446 (Fig. 1f). Immunoblotting showed higher PIEZO1 protein in both resistant lines (Fig. 1h). All nine external resistant-minus-control expression contrasts plotted in Fig. 1g were positive. The three H69AR–H69 comparisons ranged from 3.19 to 4.37 log_2_ units, whereas estimates in the H69CPR and H526 derivatives ranged from 0.25 to 1.10. In two circulating tumour cell-derived xenograft (CDX) models, PIEZO1 expression was higher at cisplatin relapse than in vehicle controls. Cisplatin IC_50_ increased from 2.32 to 15.65 µg/mL in the H69 pair and from 4.80 to 16.13 µg/mL in the H446 pair (Fig. 1i and Fig. S1g). Both derivatives also had higher etoposide IC_50_ values than their parental lines (Fig. S1h).

### PIEZO1 marks a low-neuroendocrine state across patient tumours and tumour cells

In 107 TU-SCLC tumours [9], PIEZO1 RNA correlated inversely with the NE score (Spearman ρ = −0.57, 95% confidence interval [CI] −0.69 to −0.43). The inverse relationship extended to six measurements drawn from bulk RNA, protein and tumour-cell pseudobulk RNA. The correlation with YAP induction or YAP1 protein was positive in all six measurements and ranged from 0.21 to 0.75 (Fig. 2a,b). Mouse non-NE SCLC lines expressed more Piezo1 than NE lines, and the inverse association with NE score extended to 57 human SCLC cell lines (Fig. 2c and Fig. S2a). The DepMap expression series identifies both parental lines in this context (Fig. S2d). Adjacent lung tissue contained more PIEZO1 RNA and protein than paired tumours in most patients. Accounting for tissue composition reduced those paired differences (Fig. S2b,c).

**Figure 2.**
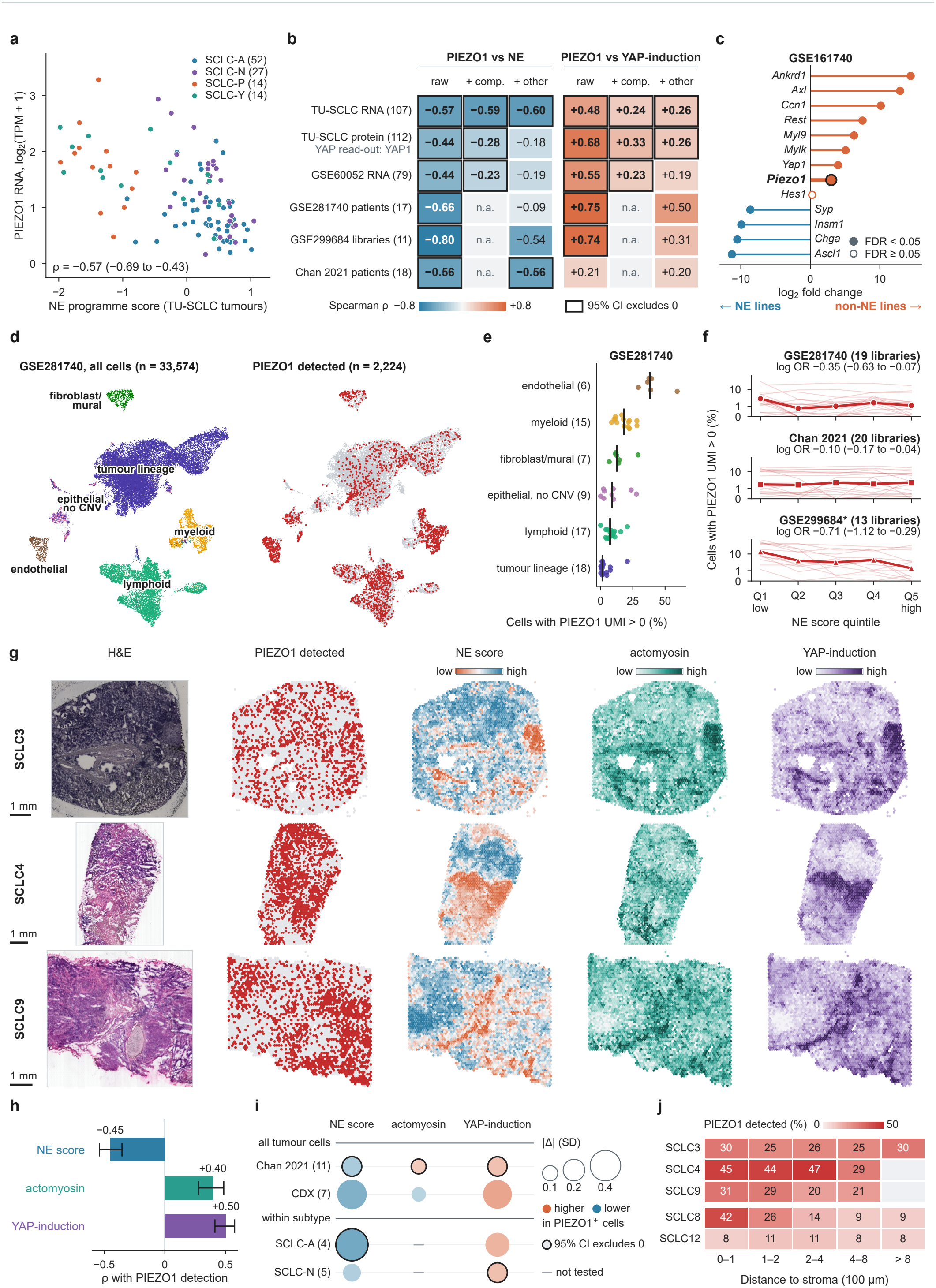
PIEZO1 expression across neuroendocrine states in SCLC. **a**, PIEZO1 RNA and NE score in 107 TU-SCLC tumours, coloured by subtype. **b**, Raw and adjusted associations of PIEZO1 with the NE and YAP programmes across six measurements. YAP1 abundance is the protein readout. **c**, Expression differences between non-NE and NE mouse SCLC lines. **d**,**e**, Cell types, PIEZO1 detection and compartment-specific detection frequencies in GSE281740. **f**, Within-library detection across NE quintiles in three human sources. Thin lines denote libraries, thick lines their medians. Pooled log ORs are per SD of NE score. The asterisk indicates patient grouping inferred from library names. **g**,**h**, Registered histology and programme maps in SCLC3, SCLC4 and SCLC9 and mean spatial correlations in tumour-rich spots. Scale bars, 1 mm. **i**, Adjusted programme differences between detected and undetected cells. Rings mark 95% CIs excluding zero. **j**, Detection frequency by distance from the stromal boundary across five sections. Empty bins contain fewer than 20 eligible spots.

Cell identity resolved this apparent discrepancy. In GSE281740, 2,224 of 33,574 cells had detectable PIEZO1. The median detection rate was 38% in endothelium and 1.5% in the tumour lineage (Fig. 2d,e). Within tumour-lineage cells, each standard deviation (SD) increase in NE score reduced the log-odds of PIEZO1 detection by 0.35 in GSE281740, 0.10 in the Chan atlas [10] and 0.71 in GSE299684 after adjustment for transcript depth and hypoxia (Fig. 2f and Fig. S3). In three tumour-rich Visium sections, spatial correlations with PIEZO1 detection averaged −0.45 for NE, +0.40 for actomyosin and +0.50 for YAP induction (Fig. 2g,h). Comparisons within tumour-cell populations placed PIEZO1-detected cells toward lower NE and higher YAP-induction activity (Fig. 2i). The stromal boundary also marked a gradient in some sections. In SCLC8, detection fell from 42% within 100 µm of the boundary to 9% beyond 800 µm, whereas SCLC3 and SCLC12 showed little variation (Fig. 2j).

### PIEZO1 detection tracks cell state and endothelial proximity

In the SC16 CDX model, PIEZO1 detection increased along a transcriptional ordering rooted in NE-high cells. The odds of detection at the low-NE end relative to the high-NE end were 1.9 in SC16, 7.3 in SC4 and 3.6 in SC53 (Fig. 3a–c). Along the SC16 ordering, ASCL1 and INSM1 detection fell as REST and YAP1 detection rose (Fig. 3d). Across CDX models, each SD increase in NE score corresponded to a change of −0.14 in the log-odds of PIEZO1 detection (95% CI −0.25 to −0.03, Fig. 3e) [11]. Inferred copy-number subclones explained 0.01–3.5% of NE-score variance in the four models that recovered simulated copy losses (Fig. S4).

**Figure 3.**
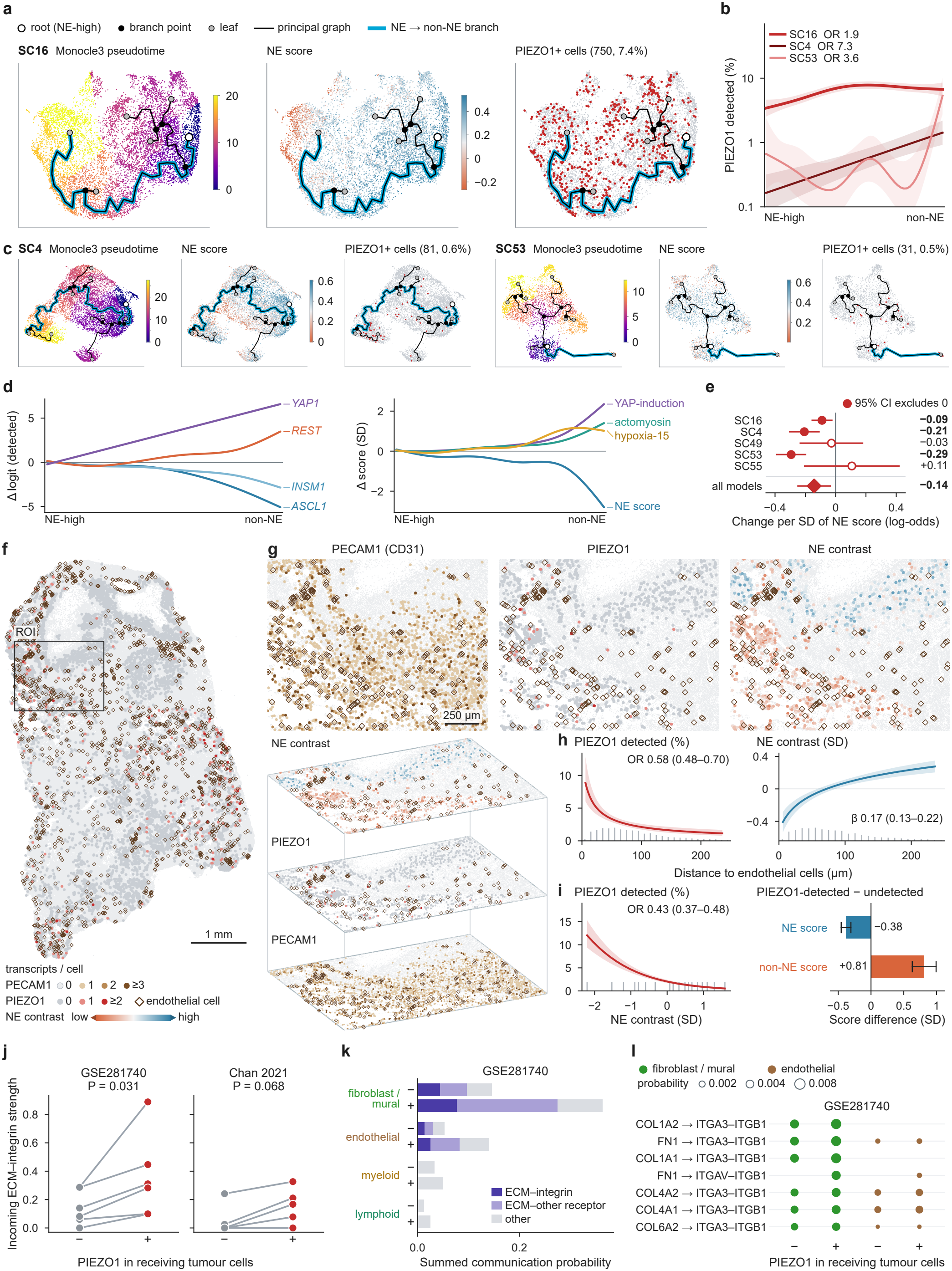
PIEZO1 along cell-state trajectories and spatial gradients. **a-d**, Monocle3 state orderings for SC16, SC4 and SC53, fitted PIEZO1 detection and changes in SC16 markers and programmes. **e**, Model-specific and pooled changes in the log-odds of detection per SD of NE score, **f**, Section p3, showing epithelial PIEZO1 counts and the selected field (scale bar, 1 mm), **g**, Co-registered PECAM1 (CD31) across all quality-controlled cells, epithelial PIEZO1 and epithelial NE contrast in the 1,600 x 1,250 pm field (scale bar, 250 μm). Diamonds mark marker-defined endothelium. The stacked view preserves tissue coordinates with schematic vertical spacing, h. Adjusted associations with distance to the nearest endothelial-cell centroid, per doubling of (distance + 10 μm), **i**. Same-cell detection odds by NE contrastand adjusted component differences. Section p3 contained 7,101 epithelial candidates, including 273 with detectable PIEZO1. Panel g shows one field. Shading and error bars in h,i denote 95% spatial-duster Cis. **j-I**. Paired incoming ECM-integrin scores, sender contributions and leading inferred ligand-receptor pairs.

To resolve epithelial and endothelial signals within the same section, we analysed six untreated primary SCLC specimens profiled by Xenium. In section p3, 273 of 7,101 epithelial candidates had detectable PIEZO1. Co-registered maps placed the epithelial signal and a lower NE contrast near annotated endothelial cells (Fig. 3f,g and Fig. S6a). For each doubling of (endothelial distance + 10 µm), the adjusted odds of PIEZO1 detection fell (odds ratio [OR] 0.583, 95% CI 0.483–0.703) and the epithelial NE contrast increased by 0.173 SD (0.128–0.219, Fig. 3h). The PIEZO1 distance association passed correction across six sections in p3. The estimates in p1 and p6 pointed in the same direction (ORs 0.830 and 0.754, adjusted q values 0.073 and 0.072, Fig. S6b–e).

The two gradients intersected in individual epithelial cells. At comparable endothelial distance, transcript depth, local cell density and lineage composition, the odds of PIEZO1 detection fell with increasing NE contrast (OR 0.425 per SD, 95% CI 0.373–0.484). Detected cells had lower NE scores (−0.381 SD) and higher non-NE scores (+0.812 SD) than undetected cells (Fig. 3i). The sign of the association persisted when each of the 14 panel genes was omitted in turn. Requiring the PIEZO1 transcript to lie at least 1 µm inside its assigned cell also preserved the association (area-adjusted OR 0.459, 95% CI 0.397–0.531, Fig. S6f–h).

In expression-based ligand–receptor analyses, predicted extracellular matrix (ECM)–integrin input was greater in PIEZO1-detected tumour cells in six of six GSE281740 samples and four of five Chan samples (paired P = 0.031 and 0.068, respectively). Fibroblast/mural and endothelial populations contributed collagen- and fibronectin-associated interactions (Fig. 3j–l). Spot-level distances and receptor detection are shown in Fig. S5.

### PIEZO1 depletion sensitizes resistant cells to cisplatin

Three siRNAs reduced PIEZO1 expression to 0.33–0.53 of non-targeting control levels across the resistant lines (Fig. 4a). With the siRNA selected for functional experiments, the cisplatin IC_50_ fell from 15.7 to 6.4 µg/mL in H69AR and from 15.7 to 5.9 µg/mL in H446DDP. In the parental lines, the PIEZO1 agonist Yoda1 [12] increased the IC_50_ from 5.2 to 9.4 µg/mL in H69 and from 6.2 to 9.7 µg/mL in H446 (Fig. 4b). Drug-free Yoda1 viability measurements across four concentrations and three time points are shown in Fig. 4c.

**Figure 4.**
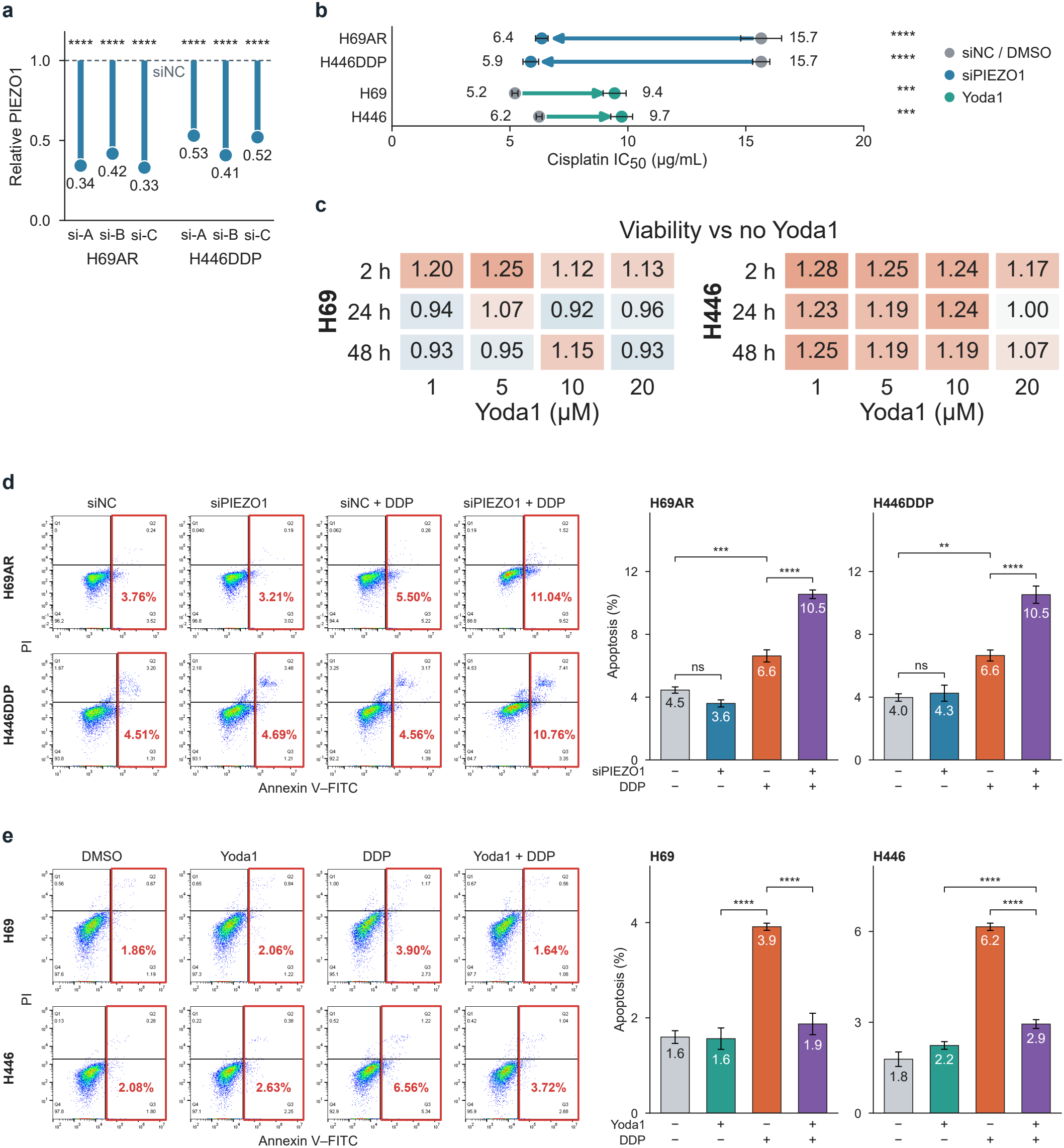
PIEZO1 perturbation alters the response to cisplatin. **a**, Relative PIEZO1 expression after three siRNAs in H69AR and H446DDP. Functional assays used one selected siRNA. **b**, Cisplatin IC_50_ following PIEZO1 knockdown in resistant lines or Yoda1 exposure in parental lines. **c**, Viability relative to matched controls without Yoda1 after Yoda1 alone. Columns show concentration and rows show exposure time. **d**, Annexin V–FITC/propidium iodide flow cytometry and apoptosis in resistant cells treated with non-targeting or PIEZO1 siRNA, with and without cisplatin. **e**, Corresponding comparisons in parental cells treated with vehicle or Yoda1. Numbers on representative cytometry plots refer to the displayed sample. Group summaries are shown alongside. DDP, cisplatin. siNC, non-targeting siRNA.

Flow cytometry linked the change in drug response to apoptosis. Under cisplatin challenge, PIEZO1 depletion increased the apoptotic fraction from 6.6% to 10.5% in both resistant lines. Yoda1 reduced cisplatin-induced apoptosis from 3.9% to 1.9% in H69 and from 6.2% to 2.9% in H446 (Fig. 4d,e). Thus, reducing PIEZO1 expression sensitized two resistant models, and agonist exposure protected both parental lines.

### PIEZO1 covaries with myosin phosphorylation and alters intracellular calcium

In 112 TU-SCLC tumours, PIEZO1 protein correlated positively with regulatory myosin light-chain phosphorylation at S19 and T18 (ρ = 0.56 and 0.59). After adjustment for light-chain family abundance and tissue composition, the corresponding partial correlations were 0.25 and 0.28. The T134 signal was weaker (partial ρ = 0.07, Fig. 5a and Fig. S2e–g). The T18/S19 peptide is shared by MYL12A, MYL12B and MYL9.

**Figure 5.**
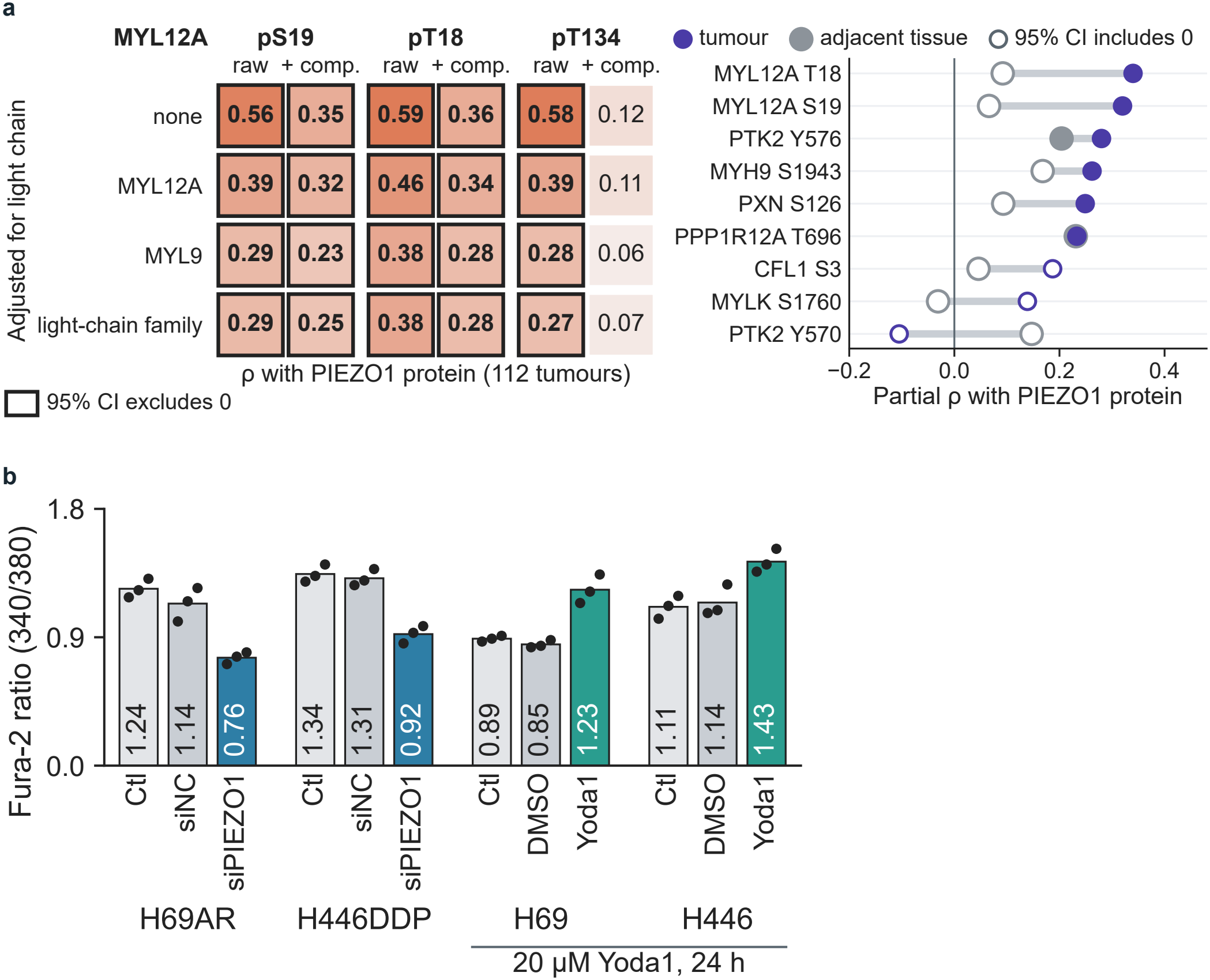
Actomyosin phosphosites and cellular calcium in relation to PIEZO1. **a**, Association of PIEZO1 protein with myosin light-chain phosphorylation in 112 TU-SCLC tumours. Left, Spearman correlations with T18, S19 and T134 under light-chain and tissue-composition adjustments. Right, adjusted phosphosite associations in tumour and adjacent tissue. Filled markers denote 95% CIs excluding zero. **b**, Fura-2 excitation ratios after PIEZO1 knockdown in H69AR and H446DDP and after 20 µM Yoda1 for 24 h in H69 and H446. Ratios compare excitation at 340 and 380 nm with the same emission wavelength.

In resistant cells, PIEZO1 depletion lowered the Fura-2 excitation ratio from 1.14 to 0.76 in H69AR and from 1.31 to 0.92 in H446DDP. In parental cells, Yoda1 raised the ratio from 0.85 to 1.23 in H69 and from 1.14 to 1.43 in H446 (Fig. 5b). Genetic depletion and agonist treatment therefore shifted the steady-state calcium readout in opposite directions.

### Yoda1 alters YAP1 localization and verteporfin attenuates its protective effect

In TU-SCLC tumours, PIEZO1 and YAP1 protein abundance were positively correlated (ρ = 0.68, 95% CI 0.56–0.77). The association persisted after adjustment for five non-tumour compartments (partial ρ = 0.33, Fig. 6a). In a published mouse SCLC model [5], induction of active YAP raised Piezo1 expression by 2.98 log_2_ units at day 21 (false discovery rate [FDR] 6 × 10^−7^), accompanied by increases in YAP target genes and a decline in NE markers (Fig. 6b).

**Figure 6.**
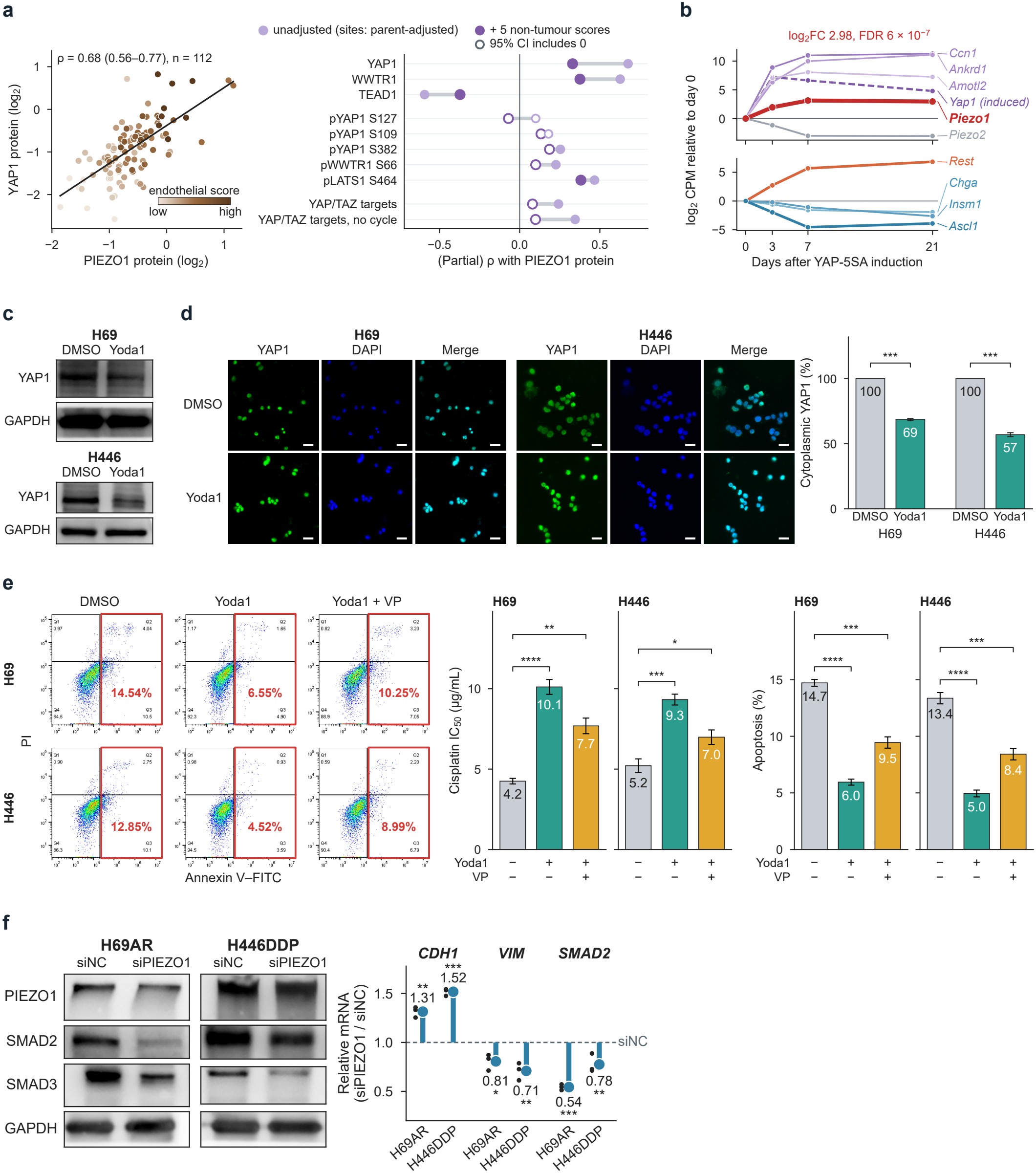
PIEZO1, YAP1 and cisplatin response in SCLC. **a**, PIEZO1 and YAP1 protein abundance in 112 tumours and associations with Hippo pathway components before and after adjustment for tissue composition. **b**, Gene expression after induction of active YAP in mouse SCLC cells (days 0, 3, 7 and 21; GSE161740). **c**,**d**, YAP1 immunoblots, immunofluorescence and cytoplasmic YAP1 signal relative to DMSO (100%) in parental lines treated with vehicle or Yoda1. Scale bars, 10 µm. **e**, Cisplatin IC_50_, representative annexin V/propidium iodide plots and apoptosis under DMSO, Yoda1 or Yoda1 plus verteporfin (VP). **f**, PIEZO1, SMAD2 and SMAD3 immunoblots with GAPDH controls, and CDH1, VIM and SMAD2 transcript levels after PIEZO1 knockdown in H69AR and H446DDP.

In parental H69 and H446 cells, Yoda1 reduced the cytoplasmic YAP1 signal to 69% and 57% of vehicle control, respectively (Fig. 6c,d). Verteporfin partly reversed Yoda1-associated protection. Adding verteporfin to Yoda1 lowered the cisplatin IC_50_ from 10.1 to 7.7 µg/mL in H69 and from 9.3 to 7.0 µg/mL in H446. Apoptosis rose from 6.0% to 9.5% and from 5.0% to 8.4%, respectively (Fig. 6e). In the resistant lines, PIEZO1 depletion also increased CDH1 and reduced VIM and SMAD2 transcripts (Fig. 6f). These responses connect PIEZO1 to a mesenchymal expression programme and place YAP-associated signalling in the response to cisplatin.

## Discussion

PIEZO1 identifies a spatially organized cell state in SCLC. Patient tumours with lower NE activity expressed more PIEZO1, an association that persisted within tumour-cell lineages. In section p3, PIEZO1-detected epithelial cells were more frequent near endothelium and had a lower NE contrast than other epithelial candidates at comparable endothelial distances. Separating the two components of the contrast showed lower NE and higher non-NE expression in the same population. This distinction matters because endothelial cells themselves had the highest frequency of PIEZO1 detection.

The perturbation experiments establish a functional role for PIEZO1 in platinum response. Depleting PIEZO1 lowered the cisplatin concentration needed to inhibit resistant cells and increased apoptosis during challenge. The effect extended across a doxorubicin-selected line and a cisplatin-selected line. Yoda1 moved the parental cells in the opposite direction, altered their calcium readout and redistributed YAP1. Verteporfin attenuated protection, consistent with a YAP-associated component. The increase in Piezo1 after YAP induction in a separate mouse model also raises the possibility of reciprocal regulation.

The spatial and functional data define a testable link between the tissue neighbourhood and treatment response. Endothelial proximity measures tissue position. The cellular assays measure drug response under defined perturbations. Establishing whether a local physical cue generates this phenotype will require measuring tissue mechanics at the mapped locations and following the response of matched epithelial cells. Spatial effects varied among the six specimens, and the cellular-resolution evidence for both gradients comes from p3. Likewise, low parental input coverage makes regional m^6^A enrichment a route to candidate discovery rather than proof of a change in site-specific methylation. The present data identify where to look for PIEZO1-expressing cells in SCLC and establish why their cisplatin response merits investigation.

## Methods

### Cell culture and perturbations

NCI-H69, NCI-H69AR and NCI-H446 cells were obtained from the American Type Culture Collection. H446DDP is a laboratory-derived cisplatin-resistant derivative of H446 [8]. Cells were cultured in RPMI 1640 at 37 °C with 5% CO_2_ and fetal bovine serum (20% for H69AR and 10% for the other lines). In cultures containing suspended and adherent populations, both fractions contributed to the assays. Three PIEZO1-targeting siRNAs (KeyGEN BioTECH) were screened against non-targeting siRNA using PepMute transfection. One selected siRNA was used in functional assays. Parental cultures received Yoda1 (20 µM for 24 h) or dimethyl sulfoxide (DMSO) vehicle [12]. Yoda1-only viability was measured at 1, 5, 10 and 20 µM after 2, 24 and 48 h. Verteporfin was used in the combination experiments shown in Fig. 6e [13].

### Viability and apoptosis

For cell counting kit-8 assays, 2 × 10^4^ cells per well were exposed to cisplatin for 24 h. The concentration series used for perturbation assays was 0, 1.5625, 3.125, 6.25, 12.5, 25, 50 and 100 µg/mL. Absorbance was read at 450 nm. Parental dose series are given in Fig. S1g. IC_50_ denotes the concentration giving 50% inhibition. Dose–response curves were fitted in GraphPad Prism. The resistance index was calculated as resistant-line IC_50_ divided by parental-line IC_50_. The H69AR control estimate in Fig. 1i also serves as the siRNA control in Fig. 4b. For apoptosis assays, cultures were challenged with cisplatin at their line-specific IC30 for 24 h and resuspended in 400 µL binding buffer. Cells were incubated in the dark at 4 °C with 5 µL annexin V–FITC for 15 min and then 5 µL propidium iodide for 5 min (Thermo Fisher Scientific) before immediate flow cytometry. Representative plots are paired with group summaries in Figs. 4d,e and 6e.

### RNA, immunoblotting, YAP1 imaging and calcium

RNA was extracted with TRIzol, reverse-transcribed with FastKing KR116 and assayed by quantitative PCR with Talent SYBR FP209. Relative expression was calculated by the 2−ΔΔCt method with GAPDH as reference. For immunoblotting, cells were lysed in RIPA buffer. Total protein was quantified by bicinchoninic acid assay, separated by SDS–PAGE, transferred to PVDF membranes and detected by chemiluminescence. Immunofluorescence images of YAP1 in parental cells were acquired with DAPI counterstaining. Cytoplasmic YAP1 signal was expressed relative to the DMSO condition within each line. For the calcium assay, cells were loaded with 1 µM Fura-2 AM for 20 min at 37 °C, washed and allowed to recover for 1 h at 37 °C in HEPES buffer. Fluorescence emission at 510 nm was measured with alternating 340- and 380-nm excitation [14]. Results are expressed as the excitation ratio.

### RNA sequencing and m^6^A enrichment

Three H69 and three H69AR RNA libraries were analysed with edgeR 4.8.2 [15]. Genes passing filterByExpr were normalized by the trimmed mean of M values. A negative-binomial quasi-likelihood model compared the two groups. Differential expression was defined by FDR < 0.05 and absolute log_2_ fold change > 1. Principal components were calculated from the 2,000 most variable genes. Hallmark enrichment used MSigDB, gene sets of 15–500 members and a signed square-root F ranking [16]. Fixed programmes were evaluated by camera [17]. The NE-up and non-NE programmes followed Zhang et al. [18], excluding YAP1, CCN1, TACSTD2 and SEZ6. The 340-gene YAP-induction programme was derived from GSE161740 after excluding Yap1 and Piezo1 and mapping one-to-one orthologues. Removing shared NE members left 143 genes. Additional fixed programmes comprised 37 published YAP/TAZ targets and 27 actomyosin genes. PIEZO1 was excluded when testing association with its expression.

For the joint screen, the provider’s differential m^6^A-enrichment estimate for the region with the largest absolute effect per gene was plotted against RNA log_2_ fold change. Regional enrichment was assessed separately using coverage reconstructed from scaled bigWigs and matched input counts. Coverage-equivalent counts were calculated as integrated covered bases divided by 200. Protein-coding genes with mean pooled input FPKM ≥1 were tiled into 100-nucleotide intervals. Adjacent tiles meeting pooled coverage and enrichment criteria were merged with condition labels masked. Nine provider regions and three independently defined regions on PIEZO1 transcript ENST00000301015 were inspected across the six libraries (Fig. S1e,f).

### External expression and patient proteogenomic data

Nine resistant-versus-control contrasts were analysed on the scale of each source. Raw counts used TMM, voom and limma [15, 19, 20]. TPM, FPKM and normalized counts used the documented log transform, and previously logged matrices retained their scale. Differences are expressed as resistant minus control in log_2_ units. Intervals were estimated where biological replicates permitted.

GSE161740 included three mouse tumour-derived lines per baseline state. The YAP induction series used two libraries per time point with a shared day-0 reference [5]. Human cell lines were drawn from DepMap 24Q4, excluding SMARCA4-undifferentiated thoracic models, and the NCI-DTP SCLC screen [21].

The TU-SCLC study provided 112 untreated surgical tumour–adjacent tissue pairs, including 107 RNA pairs [9]. GSE60052 contributed normalized expression for 79 tumours and seven normal-lung samples [22]. Within-platform associations used Spearman or partial rank correlations. In paired tissue analyses, within-patient PIEZO1 differences were regressed on differences in endothelial, immune, fibroblast, smooth-muscle and alveolar scores, with HC3 intervals. Tumour-cell pseudobulks were evaluated per patient or library. Phosphosite associations were adjusted for parent protein and tissue composition using observed measurements. Because the T18/S19 peptide maps to three regulatory myosin light-chain proteins, we also adjusted for individual light chains and for the family.

### Single-cell state scores and CDX trajectories

GSE281740 and GSE299684 cells required at least 1,000 unique molecular identifiers (UMIs), 500 detected genes and less than 20% mitochondrial reads. Epithelial assignments were checked against relative copy-number profiles with non-epithelial reference cells. Harmony was used to display the full GSE281740 atlas [23]. Chan atlas analyses used author-annotated SCLC epithelial cells and excluded RU1108 [10]. GSE281740 and Chan share eight patients and were analysed separately. The direct NE analyses included 72,875 CDX cells (21 libraries, eight models), 22,251 GSE281740 cells (19 libraries), 17,230 GSE299684 cells (13 libraries) and 41,141 Chan cells (20 batches). In GSE299684, patient groups were inferred from library prefixes.

Counts were scaled to 10,000 per cell, log-transformed after adding one, then standardized gene by gene within each library. The NE contrast was the mean of the 24 NE-up genes minus the mean of the 22 non-NE genes, using available members with nonzero variance. PIEZO1 detection required at least one raw UMI. Within libraries with ≥20 detected and ≥20 undetected cells, logistic models estimated the change in detection per SD of NE score, adjusting for transcript depth and a 15-gene hypoxia score.

Coefficients were combined within model or patient and then summarized by source using DerSimonian–Laird random effects. Expression intensity among detected cells was estimated separately. Programme differences between detected and undetected cells used depth adjustment and Hartung–Knapp intervals.

Relative copy-number profiles were inferred in eight CDX models with infercnvpy 0.4.5 (100-gene windows, 10-gene steps). Fixed state genes and the top 5% of state-correlated genes were excluded. Simulated three-to-two-copy losses in 30% of cells tested recovery, with a Jaccard cutoff of 0.5. Transcriptional orderings used untreated or vehicle libraries and 2,000 variable genes after exclusion of PIEZO1 and cell-cycle genes. Monocle3 graphs were rooted in high-NE cells, and detection along the branch ending at the lowest mean NE score was modelled with binomial generalized additive models adjusted for log UMI and library. End-to-end ORs compare the 95th and 5th percentiles of ordering rank. Diffusion pseudotime and Slingshot provided alternative orderings [24, 25, 26].

### ECM communication and multicellular spatial measurements

CellChat mass-action scores were calculated using LIANA+ 1.4.0 and CellChatDB [27, 28]. Detected and undetected tumour cells formed separate receiver groups. ECM–integrin pairs required an ECM ligand and a receptor containing an ITGA or ITGB component. Pooled analyses used 1,000 label permutations and up to 5,000 cells per group. Sample-level analyses used 300 permutations and required at least 100 tumour cells and 20 fibroblast or endothelial cells. Paired receiver contrasts required ≥10 PIEZO1-detected tumour cells.

Five untreated Visium specimens from GSE263196 were deconvolved with RCTD in full mode, using GSE281740 reference classes capped at 3,000 cells per class [29]. Three sections with ≥50 PIEZO1-detected tumour-dominant spots met the coverage rule for programme maps. Core analyses used tumour weight ≥0.7 and ≥500 UMIs after exclusion of normal-lung-dominant spots. Spatial correlations were calculated from neighbourhood-smoothed values and averaged across those sections. Distance analyses used all five sections and tumour weight ≥0.5. Spot-centre distances were measured to the nearest spot in the top decile of endothelial weight or to the nearest non-tumour spot (tumour weight <0.5). Bins with fewer than 20 tumour spots were omitted from the main boundary display.

### Cellular-resolution spatial measurements

For the six untreated primary SCLC Xenium sections (p1–p6, GSE343063), the gene-by-cell matrix and metadata were joined by exact barcode. Cells required ≥20 transcripts, positive area and finite coordinates. Lineages were assigned from epithelial (EPCAM, CDH1, CLDN7, CLDN18, MUC1), endothelial (PECAM1, CDH5, KDR, ENG, ERG, SOX17, PLVAP), immune, fibroblast and pericyte markers. Strict assignments required ≥2 detected markers, ≥3 total marker transcripts and a higher mean marker count than every other lineage. Endothelial cells additionally required CDH5. Distances were measured between centroids within connected tissue domains containing ≥5 endothelial cells and ≥200 strict epithelial candidates.

Epithelial PIEZO1 detection was modelled against log_2_(distance in µm + 10), transcript depth excluding PIEZO1, domain, cell density and endothelial, immune and stromal fractions within 100 µm. Inference used finite-sample-corrected cluster-robust standard errors across 200-µm spatial blocks and t-based 95% CIs. Benjamini–Hochberg correction covered the six section-specific distance tests. The p3 field shown in Fig. 3g was chosen by epithelial and endothelial cell coverage. Whole-domain models included 493 spatial clusters. The field and the stacked display retain their measured x–y coordinates.

The Xenium panel covered five NE genes (ASCL1, INSM1, KIF1A, RTN1, SYP) and nine non-NE genes (CAV2, CCND1, EMP1, EPHA2, ITGB4, MYOF, PLAU, PTGES, TGFBR2). Gene expression was standardized to broad epithelial candidates within section. The difference between mean NE and non-NE scores was then expressed in SD units of strict epithelial candidates. In p3 the detection model included this contrast and endothelial distance together. Linear models estimated separate component effects. Holm adjustment covered five same-cell tests and Benjamini–Hochberg correction covered 28 gene-level tests. Sensitivity analyses varied epithelial definitions, omitted genes individually, resampled 400-µm blocks and adjusted for cell area. An interior-transcript analysis required a PIEZO1 transcript with quality score ≥20 at least 1 µm inside the exported cell boundary.

### Statistical reporting

Cell assays were analysed in SPSS or GraphPad Prism using independent two-group t tests and one-way analysis of variance for multiple groups. For the experimental panels, asterisks denote P < 0.05 (*), P < 0.01 (**), P < 0.001 (***) and P < 0.0001 (****). Values marked ns did not reach the 0.05 threshold. Independent culture-batch counts were unavailable for some legacy assays. Sequencing, phosphoproteomic and spatial comparisons used the statistical units and multiplicity adjustments specified above.

## Data and code availability

Public RNA and spatial data are available through the Gene Expression Omnibus under GSE60052, GSE138267, GSE161740, GSE281740, GSE299684, GSE263196, GSE343063, GSE223949, GSE152529, GSE10841, GSE151904 and GSE269636. TU-SCLC proteogenomic data are associated with PRJCA012926 [9]. The Chan atlas is available through CELLxGENE (collection 62e8f058-9c37-48bc-9200-e767f318a8ec) [10]. DepMap release 24Q4 is archived under doi:10.25452/figshare.plus.27993248.v1. The H69CPR array comparison used NCI-DTP SCLC CellMinerCDB [21]. Study-generated RNA-seq and m^6^A-enrichment data and the underlying cell-assay records have not been deposited in a public repository. Custom analysis code is available from the corresponding authors upon request.

## Ethics

Patient tissue and CDX analyses used data released by the original investigators under the approvals reported in the source studies. The experiments reported here used established human SCLC cell lines.

## Author contributions

Z.W. performed the experiments and drafted and edited the manuscript. L.Z. and X.S. curated the data. L.Z. and Z.W. prepared the visualizations. Z.W. and L.Z. performed statistical analyses. F.Z. and M.L. reviewed the manuscript and administered the project.

## Funding

This work was supported by the Guangdong Basic and Applied Basic Research Foundation (2020A1515011454), the National Natural Science Foundation of China (81902311), and the First Affiliated Hospital of Guangzhou Medical University Nanshan Talents Program (2022111708151837).

## Competing interests

The authors declare no competing interests.

## Notes

### Competing Interest Statement

The authors have declared no competing interest.

### Summary of Updates

This revision adds single-cell spatial analyses of six primary small cell lung cancer specimens. In one section, epithelial PIEZO1 detection was associated with both proximity to endothelium and lower neuroendocrine gene expression in the same cells. We expanded comparisons across patient datasets and drug-resistant models, reorganized six main figures and six supplementary figures, and updated the title, abstract and methods.

