## Supplementary Information (Figures S1-S6) for "Periendothelial enrichment of PIEZO1 and its contribution to cisplatin tolerance in small cell lung cancer"

#### Supplementary Figure S1

### RNA-seq quality, gene programmes, PIEZO1 region coverage and drug responses

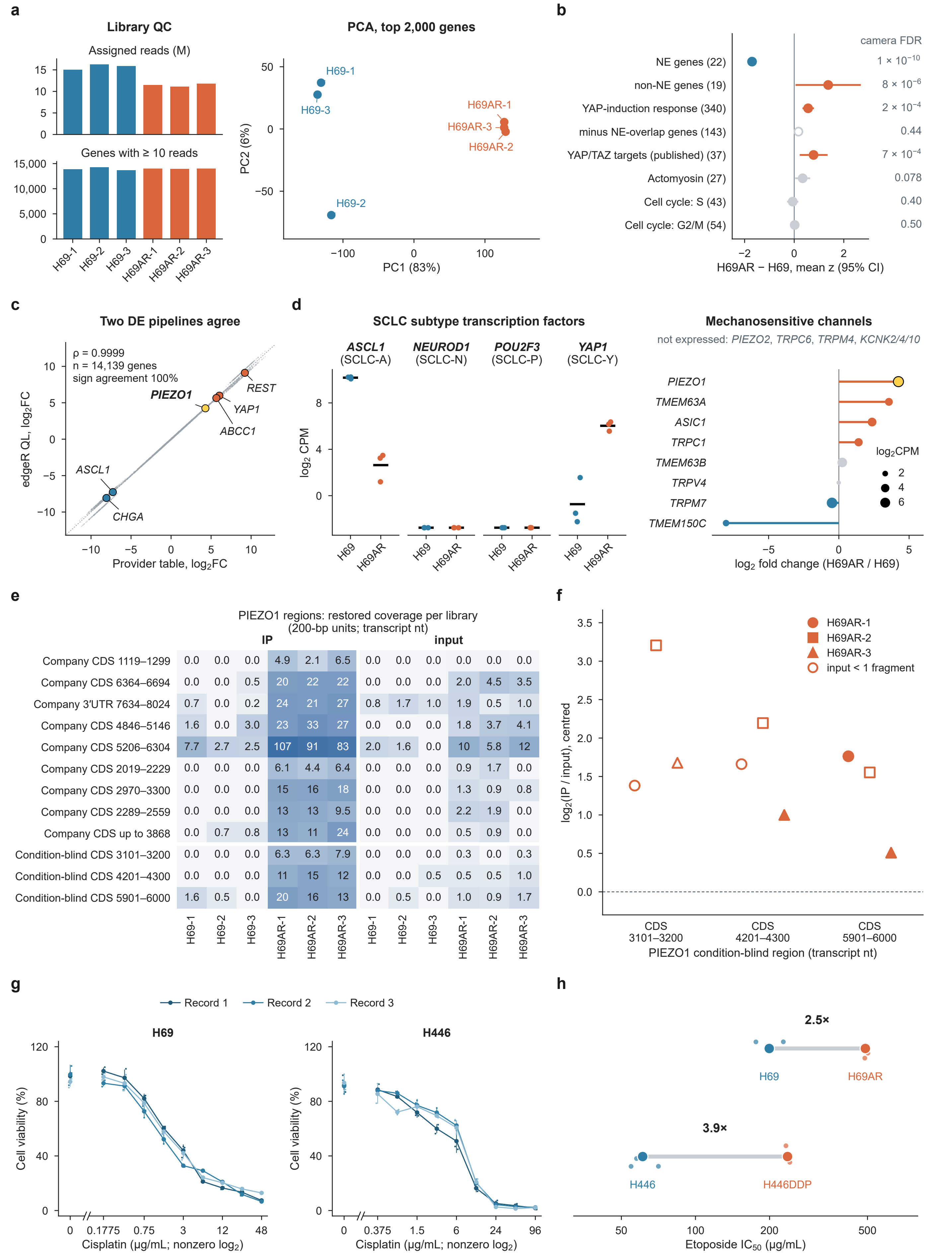


**a,** RNA library quality (assigned reads; genes with ≥10 reads) and principal-component analysis (PC1 83%, PC2 6%). **b,** Fixed programmes, H69AR − H69 (mean z, 95% CI, CAMERA FDR): NE genes (22; $1 \times 10^{-10}$), non-NE genes (19; $8 \times 10^{-6}$), YAP-induction response (340; $2 \times 10^{-4}$), the same response without its NE-overlapping genes (143; 0.44), published YAP/TAZ targets (37; $7 \times 10^{-4}$), actomyosin (27; 0.078), and cell cycle S (43; 0.4) and G2/M (54; 0.5). **c,** Provider table and local edgeR log2 fold changes (ρ = 0.9999, 14,139 genes, sign agreement 100%). **d,** SCLC subtype transcription factors (ASCL1, NEUROD1, POU2F3, YAP1; log2 CPM) and mechanosensitive channels (log2 fold change, H69AR versus H69).

**e,** Restored coverage of the nine provider-defined and three condition-blind PIEZO1 regions in every IP and input library. **f,** IP/input ratio (log2, centred) of the three condition-blind PIEZO1 regions in the three H69AR libraries. **g,** Cisplatin dose–response records of H69 and H446 (three records each). **h,** Etoposide IC50 of H69AR (2.5-fold over H69) and H446DDP (3.9-fold over H446).

NE, neuroendocrine; IP, immunoprecipitation; CPM, counts per million; CI, confidence interval; FDR, false discovery rate.

#### Supplementary Figure S2

### PIEZO1 in SCLC cell lines, paired tissues and tumour phosphoproteomes

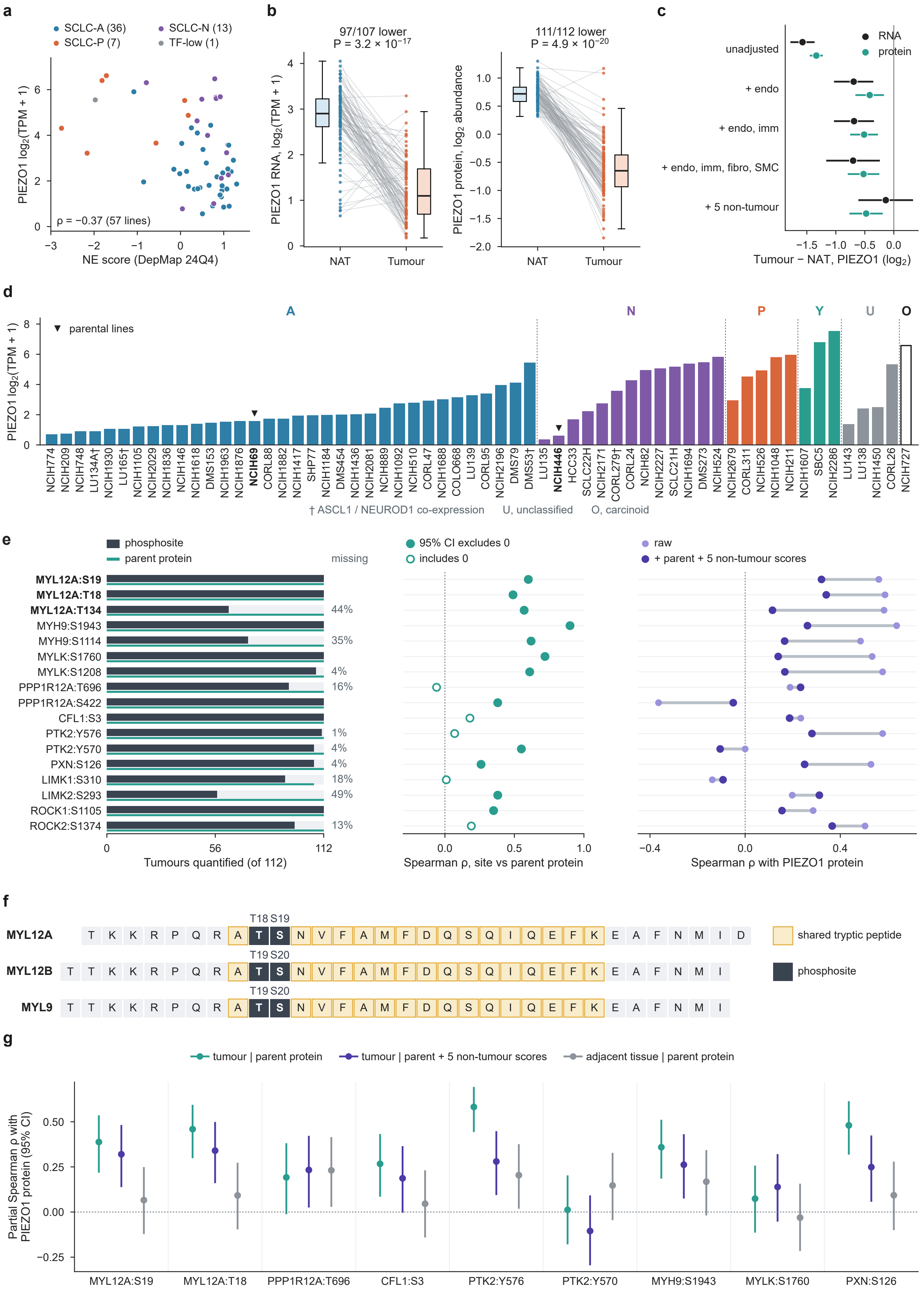


**a,** PIEZO1 expression and NE score across 57 DepMap 24Q4 SCLC lines (Spearman ρ = −0.37). **b,** PIEZO1 RNA and protein in paired TU-SCLC tumours and adjacent tissues. Tumour PIEZO1 was lower in 97 of 107 RNA pairs ($P = 3.2 \times 10^{-17}$) and 111 of 112 protein pairs ($P = 4.9 \times 10^{-20}$). **c,** Tumour-minus-adjacent differences in PIEZO1 RNA and protein after sequential adjustment for non-tumour cell content, with 95% CIs. **d,** Baseline PIEZO1 expression across 60 DepMap lines coloured by subtype, with parental H69 and H446 marked.

**e,** Coverage of 17 phosphosites and their parent proteins across 112 TU-SCLC tumours, site–parent correlations and associations between each phosphosite and PIEZO1 protein before and after adjustment. **f,** Shared tryptic peptide carrying MYL12A T18/S19, corresponding to T19/S20 in MYL12B and MYL9. **g,** Partial Spearman correlations with PIEZO1 protein at nine actomyosin or adhesion sites (95% CIs). Tumour estimates adjust for the parent protein alone or with five non-tumour scores. Adjacent-tissue estimates adjust for the parent protein.

NE, neuroendocrine; NAT, normal adjacent tissue; CI, confidence interval.

#### Supplementary Figure S3

### Single-cell sources and neuroendocrine associations across patients, models and libraries

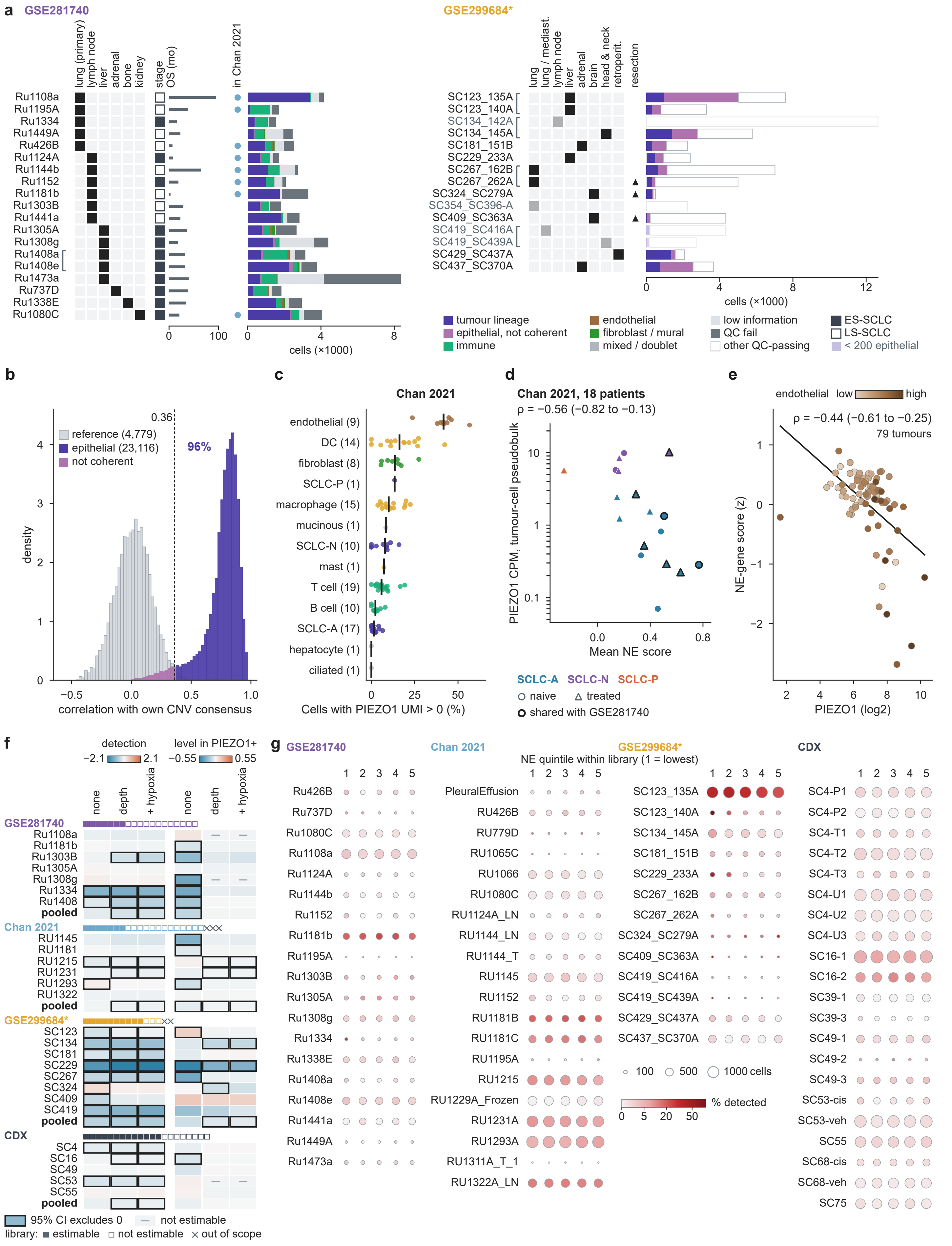


**a,** Sample information and cell composition in GSE281740 (19 samples) and GSE299684* (15 libraries). A bracket links Ru1408a and Ru1408e. The asterisk denotes patient grouping inferred from library names. **b,** Epithelial copy-number coherence relative to reference cells. Of the epithelial cells, 96% exceeded the 0.36 threshold. **c,** Sample-level PIEZO1 detection by cell type in Chan 2021. **d,** Tumour-cell pseudobulk PIEZO1 expression against mean NE score in Chan 2021 (18 patients, Spearman ρ = −0.56, 95% CI −0.82 to −0.13). **e,** PIEZO1 expression and NE-gene score in 79 GSE60052 tumours coloured by endothelial content (ρ = −0.44, 95% CI −0.61 to −0.25).

**f,** Unit-level and pooled associations per 1-SD increase in NE score for PIEZO1 detection and expression among detected cells. Columns show unadjusted, depth-adjusted and depth-plus-hypoxia-adjusted estimates. **g,** PIEZO1 detection by NE quintile in 73 libraries from GSE281740 (19), Chan 2021 (20), GSE299684* (13) and CDX (21).

NE, neuroendocrine; CNV, copy-number variation; SD, standard deviation; CI, confidence interval.

#### Supplementary Figure S4

### Copy-number subclones and cell-state ordering in CDX models

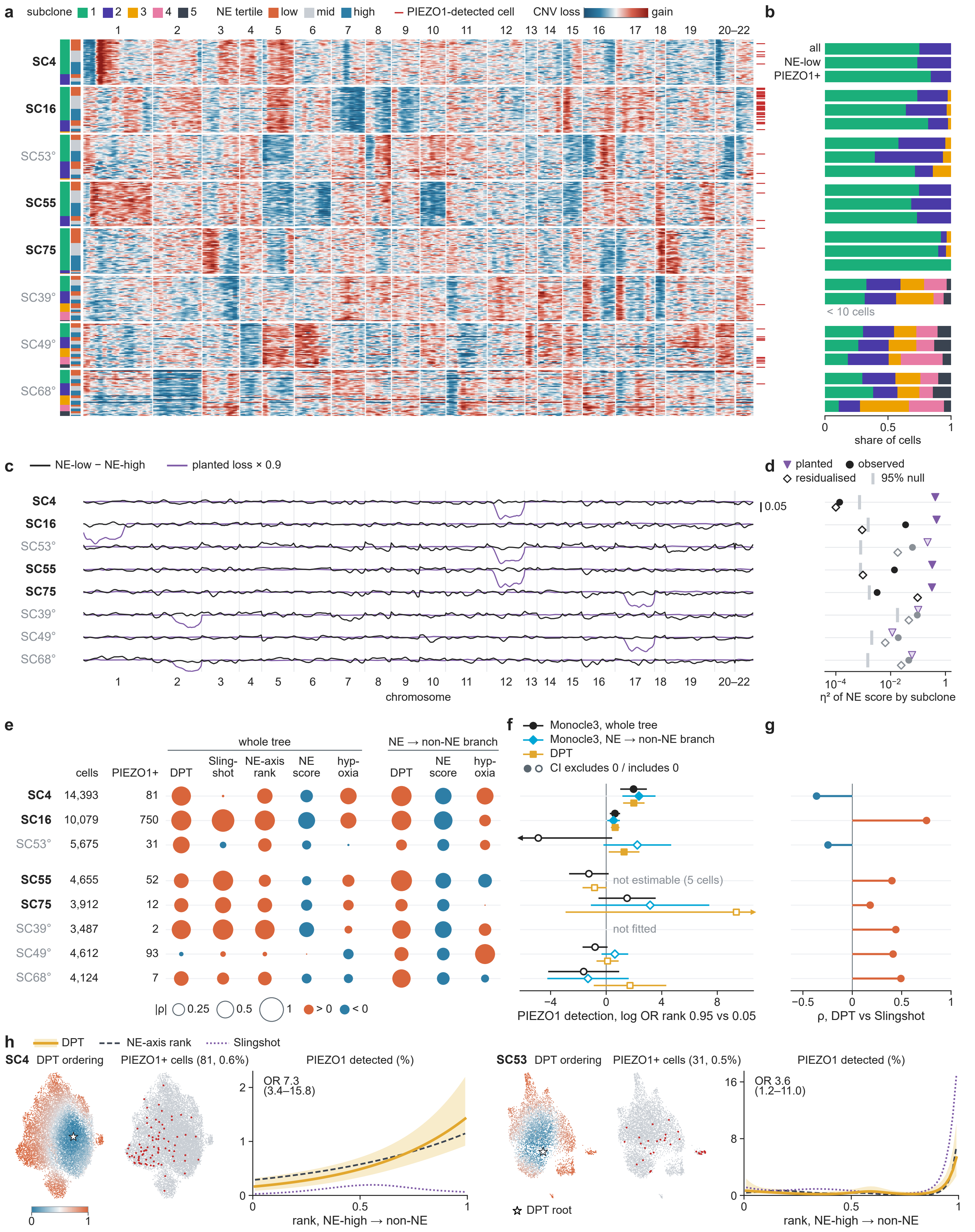


**a,** Relative CNV heat maps of the eight CDX models with subclone, NE tertile and PIEZO1-detected cells marked. **b,** Share of each CNV subclone among all cells, NE-low cells and PIEZO1-detected cells. **c,** NE-low minus NE-high CNV profile along the chromosomes, with the planted one-copy loss for scale. **d,** Share of NE-score variance explained by subclone (η²): observed, with planted control, residualized, and the 95% null.

**e,** |ρ| (circle size; orange, ρ > 0; blue, ρ < 0) of Monocle3 pseudotime with DPT, Slingshot, NE-axis rank, NE score and hypoxia score over the whole tree and with DPT, NE score and hypoxia score along the NE → non-NE branch, for each model, with cell and PIEZO1-detected counts. **f,** PIEZO1 detection along the Monocle3 whole-tree, Monocle3 NE → non-NE branch and DPT orderings, log odds ratio between rank 0.95 and 0.05, per model (SC39 not fitted; SC55 Monocle3 NE → non-NE branch not estimable, 5 cells). **g,** Correlation of DPT with Slingshot. **h,** Diffusion pseudotime maps, PIEZO1-detected cells and fitted detection curves for SC4 and SC53. End-to-end detection ORs were 7.3 (95% CI 3.4–15.8) for SC4 and 3.6 (1.2–11.0) for SC53.

CDX, circulating tumour cell-derived xenograft; CNV, copy-number variation; NE, neuroendocrine; DPT, diffusion pseudotime; OR, odds ratio; CI, confidence interval.

Supplementary Figure S5

### Spatial sections and extracellular matrix receptor detection

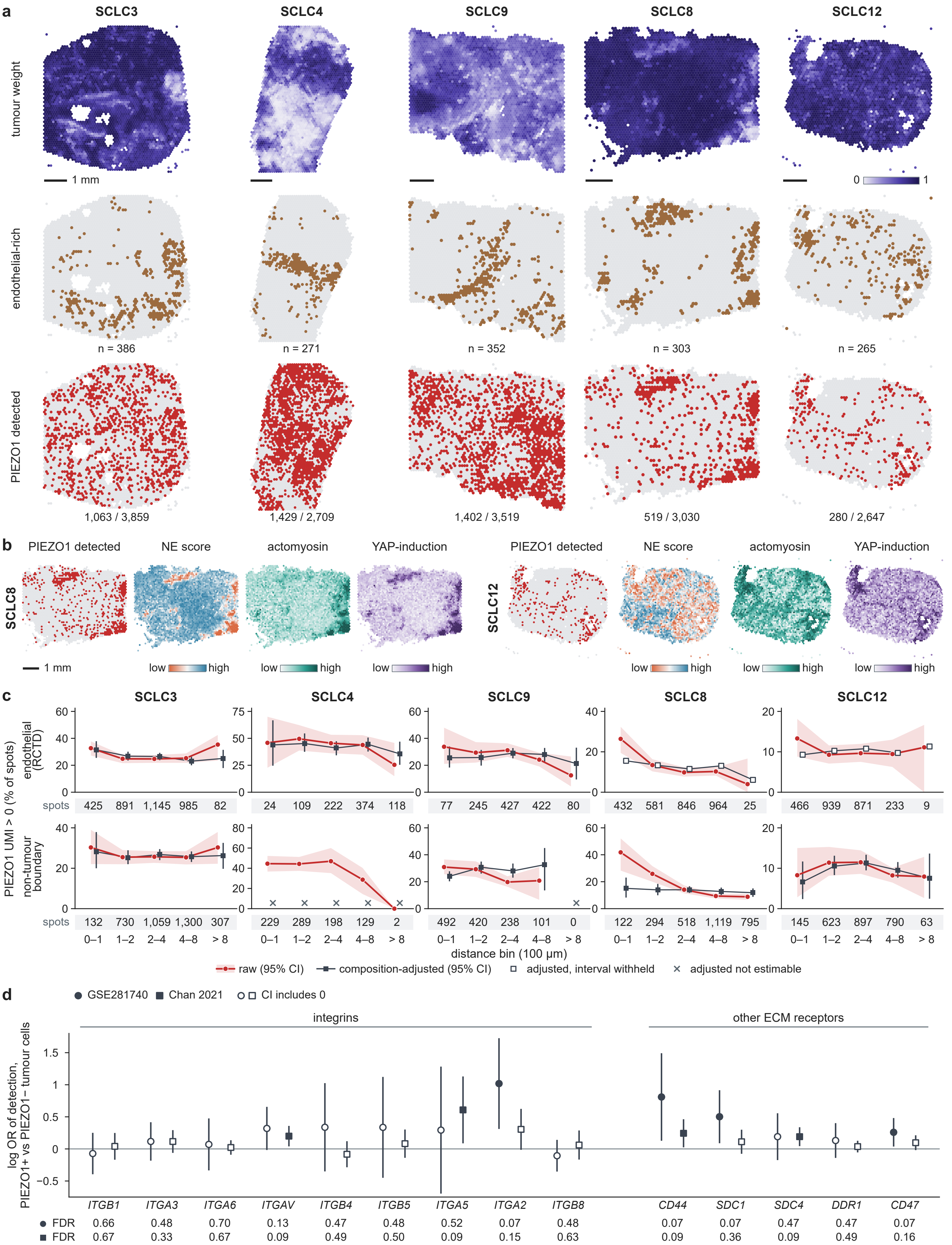


**a,** Tumour weights estimated by robust cell type decomposition, endothelial-rich spots (n = 386, 271, 352, 303, 265) and PIEZO1-detected spots (1,063 of 3,859; 1,429 of 2,709; 1,402 of 3,519; 519 of 3,030; 280 of 2,647) in SCLC3, SCLC4, SCLC9, SCLC8 and SCLC12. **b,** PIEZO1-detected spots and NE, actomyosin and YAP-induction maps of SCLC8 and SCLC12.

**c,** PIEZO1 detection by distance bin to endothelial-rich spots and to the non-tumour boundary, raw and composition-adjusted, in the five sections. **d,** Detection of nine integrins (ITGB1, ITGA3, ITGA6, ITGAV, ITGB4, ITGB5, ITGA5, ITGA2, ITGB8) and of CD44, SDC1, SDC4, DDR1 and CD47 in PIEZO1-detected versus undetected tumour cells (log odds ratio, 95% CI, FDR) in GSE281740 and Chan 2021.

NE, neuroendocrine; ECM, extracellular matrix; CI, confidence interval; FDR, false discovery rate.

#### Supplementary Figure S6

### Xenium sections and same-cell analyses

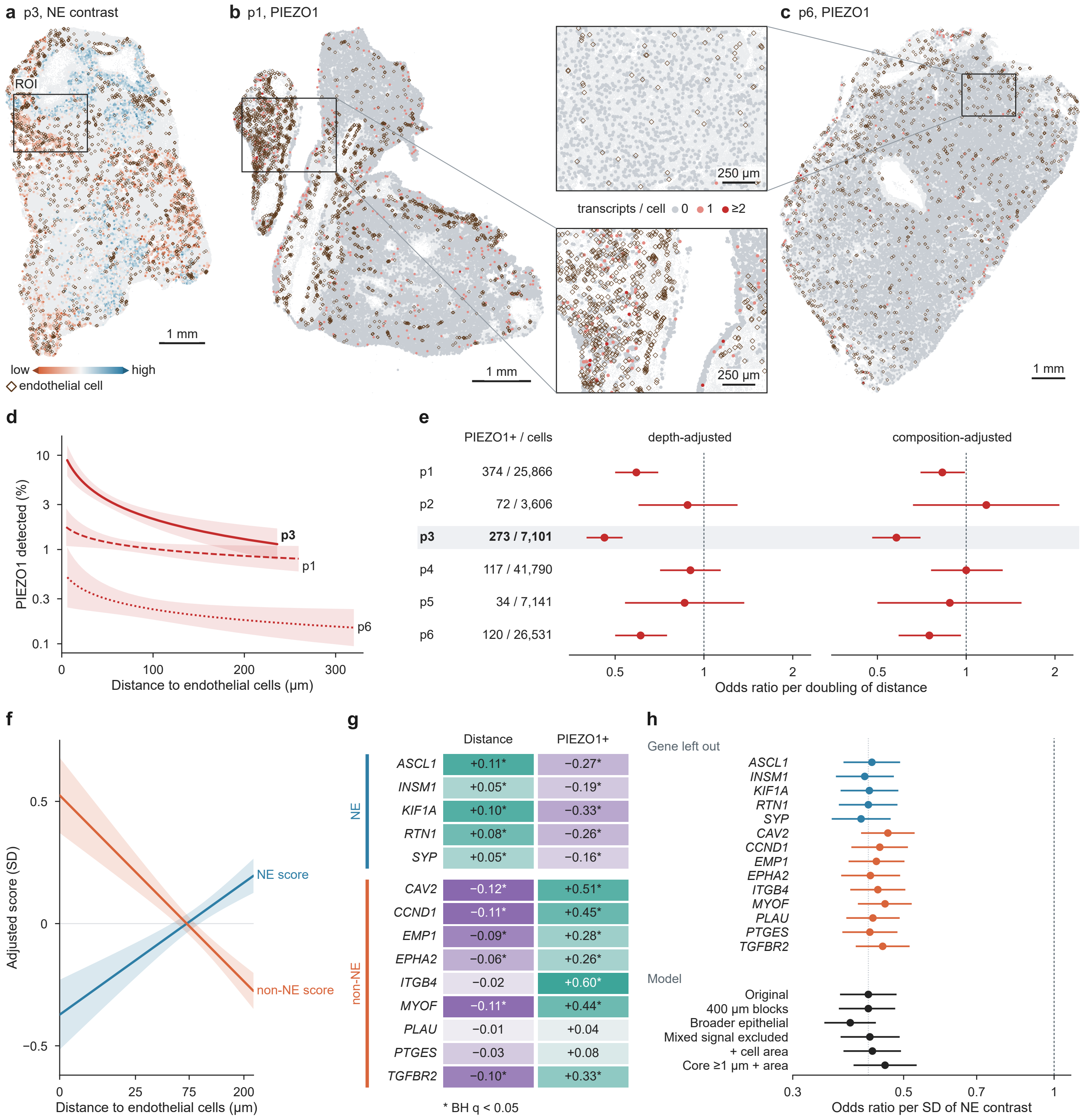


**a,** Whole-section map of epithelial NE contrast in p3. Orange denotes lower and blue higher contrast. **b,** Whole-section and enlarged maps of epithelial PIEZO1 in p1 (374 of 25,866 candidates detected). **c,** Corresponding maps in p6 (120 of 26,531 detected). **d,** Adjusted probability of detecting PIEZO1 by distance to the nearest endothelial-cell centroid in p1, p3 and p6. **e,** Depth-adjusted and composition-adjusted ORs per doubling of (distance + 10 µm) in all six sections, with 95% CIs and detected and total cell counts.

**f,** Adjusted p3 NE and non-NE scores by endothelial distance. Slopes per doubling of (distance + 10 µm) were +0.123 SD and −0.173 SD, respectively. **g,** Standardized coefficients for all 14 covered genes in p3. The first column gives the distance coefficient and the second the difference between PIEZO1-detected and undetected cells at matched covariate values. Colours are scaled within each column, with purple indicating negative and teal positive coefficients. Asterisks indicate Benjamini–Hochberg q <0.05 across 28 tests. **h,** ORs for PIEZO1 detection per 1-SD increase in NE contrast after omission of each gene (left), and under the original model and five sensitivity analyses (right). These used 400-µm spatial blocks, a broader epithelial definition, exclusion of mixed-lineage signal, adjustment for cell area, or detection of a transcript at least 1 µm inside the exported cell boundary with area adjustment. Bars show 95% intervals.

NE, neuroendocrine; SD, standard deviation; OR, odds ratio; CI, confidence interval.
